# Embryo movement is required for limb tendon maturation

**DOI:** 10.1101/2024.07.18.604105

**Authors:** Rebecca A. Rolfe, Ebru Talak Busturkmen, Lauren Sliney, Grace Hayden, Nicholas Dunne, Niamh Buckley, Helen McCarthy, Spencer E. Szczesny, Paula Murphy

## Abstract

Following early cell specification and tenocyte differentiation at the sites of future tendons, very little is known about how tendon maturation into robust load-bearing tissue is regulated. Between embryonic day (E)16 and E18 in the chick, there is a rapid change in mechanical properties which is dependent on normal embryo movement. However, the tissue, cellular and molecular changes that contribute to this transition are not well defined. Here we profiled aspects of late tendon development (collagen fibre alignment, cell organisation and Yap pathway activity), describing changes that coincide with tissue maturation. We compared effects of rigid (constant static loading) and flaccid (no loading) immobilisation to gain insight into developmental steps influenced by mechanical cues. We show that YAP signalling is active and responsive to movement in late tendon. Collagen fibre alignment increased over time and under static loading. Cells organise into end-to-end stacked columns with increased distance between adjacent columns, where collagen fibres are deposited; this organisation was lost following both types of immobilisation. We conclude that specific aspects of tendon maturation requires controlled levels of dynamic muscle-generated stimulation. Such a developmental approach to understanding how tendons are constructed will inform future work to engineer improved tensile load-bearing tissues.

## 1 Introduction

Tendons and ligaments are connective tissues within the musculoskeletal system that must sustain large mechanical loads. While ligaments connect bone to bone, tendons play a key role in integrating the musculoskeletal system, transmitting the loads produced by muscle contraction to the skeleton to facilitate movement. Despite hierarchical structures that are honed to provide the robust tensile strength required, tendons and ligaments are prone to injury and wear and tear, leading to pain and disability(Millar et al., 2021). This presents acute clinical challenges due to the limited healing capabilities of such hypocellular and hypovascular tissues, with long-term morbidity at the site, even when surgical repair is attempted. For all of these reasons tendons are prime targets for tissue engineering approaches, yet to date attempts have failed to produce structurally robust tissue with appropriate biomechanical properties for regenerative therapies. A developmental biology approach is required to understand each of the steps involved, and the mechanisms that drive natural assembly of these highly ordered and structurally important tissues in order to recapitulate or mimic the processes in vitro(Glass et al., 2014; Szczesny and Corr, 2023).

While much has been uncovered about the origin of progenitor cells and early tendon development, less is known about important steps during later development when rapid changes occur in macroscale mechanics to form a tissue capable of bearing a load(McBride, 1988). Tendon primordia are already detected in place in the early chick limb bud(Kardon, 1998; Roddy et al., 2011), and key signalling pathways and transcription factors required for early development have been identified, such as Scleraxis (Scx) and Egr1 which responds to TGFβ, FGF signalling and mechanical cues(Gaut et al., 2016; Havis et al., 2016; Schweitzer et al., 2001). The transmembrane glycoprotein Tenomodulin (Tnmd) is a regulator of cell proliferation and collagen fibril maturation(Docheva et al., 2005; Shukunami et al., 2006) while Mohawk (Mkx) is a transcription factor that also regulates tendon differentiation(Liu et al., 2015). The predominant collagen expressed is collagen 1 with minor contributions from collagens 3 and 5(Silver et al., 2003). During development tenocytes align longitudinally forming linear channels for collagen deposition and fibril formation(Bobzin et al., 2021; Kalson et al., 2010; Richardson et al., 2007). Collagen fibrils align into fibres and are bundled into fascicles forming the tendon. It is this hierarchical structure of the extracellular matrix (ECM) that gives the tendon its biomechanical properties. The ratio of cell to extracellular matrix changes as tendons mature and, at a critical timepoint (Embryonic day (E) 16-18 in the chick), there is a rapid increase in fibre stiffness and strength(Birk et al., 1995; McBride, 1988; Peterson et al., 2021). Very little is known about the cellular and molecular changes that take place and what drives these rapid changes in biomechanical properties although the transition has been shown to be dependent on embryo movement(Peterson et al., 2021).

Embryo movement is essential for normal development of multiple components of the musculoskeletal system including bone size, shape and mechanics, and joint formation(Felsenthal and Zelzer, 2017; Murphy and Rolfe, 2023). While mechanical forces from movement have been shown to be important at multiple points in tendon development(Edom-Vovard et al., 2002; Germiller et al., 1998), designing the experiments to specifically examine the critical period of transition in biomechanical properties has shown that embryonic immobilisation results in reduced mechanical modulus and failed developmental maturation(Pan et al., 2018; Peterson et al., 2021). We have previously shown that rigid paralysis leads to reduction in the size of tendons and fibril diameters(Peterson et al., 2021). Further investigation of the impact of immobilisation is needed to shed light on the cellular, molecular and biomechanical changes that are critical for tendon maturation.

Very little is known about the molecular mechanisms that interpret mechanical cues generated by movement in the skeletal system. We have previously implicated Wnt and BMP signalling as sensors that respond in the forming joint region(Rolfe et al., 2018; Singh et al., 2018) and YAP signalling in localised growth required for bone shape(Shea et al., 2020). YAP is of particular interest as it is known to respond to external environmental cues in a number of contexts ((reviewed in Panciera et al., 2017)) including cell fate decisions in the early embryo ((reviewed inYildirim et al., 2021)) and in the control of tissue structure and size in developing lung, kidney, brain epidermis, liver, bone, teeth and heart ((reviewed inDavis and Tapon, 2019; Mia and Singh, 2019; Pibiri and Simbula, 2022; Yang et al., 2023); Zhao et al., 2021). The famous work of Piccolo and colleagues demonstrated that YAP is regulated by the mechanical environment with mesenchymal stem cells grown on rigid surfaces resulted in more nuclear YAP activity and osteogenic differentiation, while softer matrices resulted in cytoplasmic YAP and adipogenesis(Dupont et al., 2011). Very little is known about YAP signalling during tendon development, apart from transcriptomic evidence of YAP target gene expression during late stages(Yeung et al., 2015), however there is evidence that YAP can influence tenogenesis in vitro; with inhibition of YAP downregulating tendon related gene expression in differentiating mesenchymal stem cells(Li et al., 2022), enhanced nuclear translocation of YAP reported to promote tenogenic differentiation(Tao et al., 2021; Tomás et al., 2019), upregulation of YAP following treatment with a novel exercise-induced myokine (Irisin) that promotes tenogenic proliferation and differentiation in culture(Xu et al., 2022) and improved performance of progenitor cells following YAP activation in culture(Lu et al., 2023). Furthermore, YAP has been shown to be critical to sensing tensional homeostasis in mature tendons(Jones et al., 2023). YAP is therefore a prime candidate to investigate as a mediator of mechanosensitive tendon maturation.

To enhance understanding of the tissue, cellular and molecular level changes that take place during tendon maturation, here we examine embryonic chick metatarsal tendons across late stages of development, quantifying changes in collagen fibre alignment, cell density, nuclear shape and YAP pathway component levels. We expand our previous analysis(Peterson et al., 2021) of the impact of immobilisation on tendon maturation by comparing rigid and flaccid paralysis across multiple tendons. We show effects of immobilisation on tendon size, spacing, collagen fibre alignment and cellular organisation. Having previously shown that YAP signalling is altered in the skeletal rudiments of muscle-less mouse embryos(Shea et al., 2020), we explored, for the first time, the potential involvement of YAP signalling in mechanoregulation of embryonic tendon maturation.

## 2 Materials and methods

### 2.1 Egg incubation and *In ovo* movement manipulation

Fertilised eggs (Ross 308, supplied by Allenwood Broiler Breeders), were incubated at 37.7°C in a humidified incubator. Work on chick embryos does not require a licence from the Irish Ministry of Health under European Legislation (Directive 2010/63/EU), all work on chick embryos was approved by the Trinity Ethics committee. Following 3 days of incubation, 5mls of albumen was removed from each egg using an 18-gauge needle. Embryos were sacrificed daily from E14 (HH40) to E20 (HH46) to assess the structural and molecular changes that occur during late tendon development.

Immobilisation treatments consisted of daily application of 0.2% Decamethonium bromide (DMB) (rigid paralysis) or 0.2% Pancuronium bromide (flaccid paralysis) (both Sigma-Aldrich) in sterile Hank’s Buffered Saline (HBSS) (Gibco) plus 1% antibiotic/ antimycotic (aa) (Penicillin, streptomycin, amphotericin B; Sigma-Aldrich), dripped directly onto the vasculature of the chorioallantoic membrane through the “windowed” egg. Immobilisation began at E16 (100 µl 0.2% DMB or 0.2% PB) with subsequent daily treatments (50 µl) until harvest. Controls were treated with equal volumes of HBSS plus aa. The experiment was repeated independently three times. Control and immobilised embryos were sacrificed at E20 (HH46) to investigate the structural and molecular effects of immobilisation.

Each egg was place on ice for at least 15 minutes prior to euthanasia by decapitation. Embryos were dissected in ice cold Phosphate Buffered Saline (PBS). Each embryo was staged using Hamburger and Hamilton criteria(Hamburger and Hamilton, 1992). Limbs were dissected and either fixed in 4% paraformaldehyde (PFA) in PBS at 4°C at least overnight, or metatarsal ankle tendons were sub-dissected and placed in TRIzol for subsequent protein and RNA extraction.

### 2.2 Histological tissue processing

Fixed hindlimbs were placed in 10% EDTA (pH 7.4) for at least 7 days at 4°C for decalcification and subsequently dehydrated through a graded series of ethanol (ETOH) (25%, 50% 70%, for a minimum of 1x 1 hour washes. Limbs were stored in 70% ETOH prior to processing for paraffin embedding or cyosectioning. One hindlimb from each specimen was processed for paraffin sectioning. Paraffin processing involved subsequent 2 x 1 hour washes in 100% ETOH, 2 x Histoclear (National Diagnostics) washes (1 hour each) and a minimum of 5 x 1hour paraffin washes prior to embedding in paraffin. A full series of longitudinal (8µm) sections through the ankle joint, and a series of cross sections (8µm) through the midpoint of the tarsometatarsal region, at a distance approximately 1-2mm proximal to digit two of the foot were prepared for each specimen. Paraffin sections were dewaxed, rehydrated, and stained to highlight connective tissue using Masson Trichrome (Sigma-Aldrich, HT15) or further processed for protein localisation (cf 2.3 and 2.4).

Hindlimbs stored in 70% EtOH processed for cryosectioning were rehydrated and cryoprotected in 30% sucrose in PBS overnight. Longitudinal sections (10 µm) were prepared for immunolabelling.

### 2.3 Picrosirius red staining and polarised light microscopy (PLM)

Collagen fibre alignment was examined in longitudinal sections of metatarsal tendons across development (E14, E16, E18 and E20) and following immobilisation. Collagen was stained with Picrosirius Red (0.1% Direct Red 80 in saturated Picric acid solution (Sigma P6744)) for 1 hour, and the samples were examined under polarised light (Olympus BX41TF Microscope fitted with polariser and analyser filters). Images were captured using Ocular Software. The sections were consistently oriented with the tendon long axis running left to right on the imaging screen and images were captured with the 20X objective, first under white light to image the ROI, followed by two polarised light images (Figure 1A). The polarised light images were selected by first setting the polariser filter to 0⁰ and rotating the analyser filter until the brightest birefringence pattern was visible (Image 1; angle 0⁰) and then moving the polariser filter to 45⁰ (Image 2; angle 45⁰). Using ImageJ, the two images were merged, converted to 8bit and a central portion of the tendon cropped to 500 x 500 µm for further analysis (Figure 1B-D). Fibre alignment analysis was performed on each ROI using the Plugin OrientationJ tool in ImageJ (Figure 1E-G)(Rezakhaniha et al., 2012), specifically using “Distribution for Histogram” and “Analysis for Colour Map” functions. This produces a colour map of orientations and allows a histogram of local orientations to be generated (Figure 1G). A minimum of 3 ROIs were measured for each independent biological sample (chick embryo) across developmental times and following treatment. Specifically, for E14 and E20 10 ROIs were analysed across each of 4 embryos; for E16 10 ROIs across each of 3 embryos; for E18, 17 ROIs across each of 5 embryos. For comparison of control and immobilised, 49 ROIs across each of 14 Control embryos, 27 ROIs across 7 Rigid immobilised embryos and 23 ROIs across 6 Flaccid immobilised embryos. Fibre alignment analysis data was represented as proportion of fibres within three orientation categories (+/-10⁰, +/-10-20⁰, +/-20-30⁰) across each group for comparison and statistical analysis. Statistical analysis was carried out by univariate ANOVA followed by Tukey post hoc tests. *p≤0.05, **p≤0.01, ***p≤0.001

**Figure 1:**
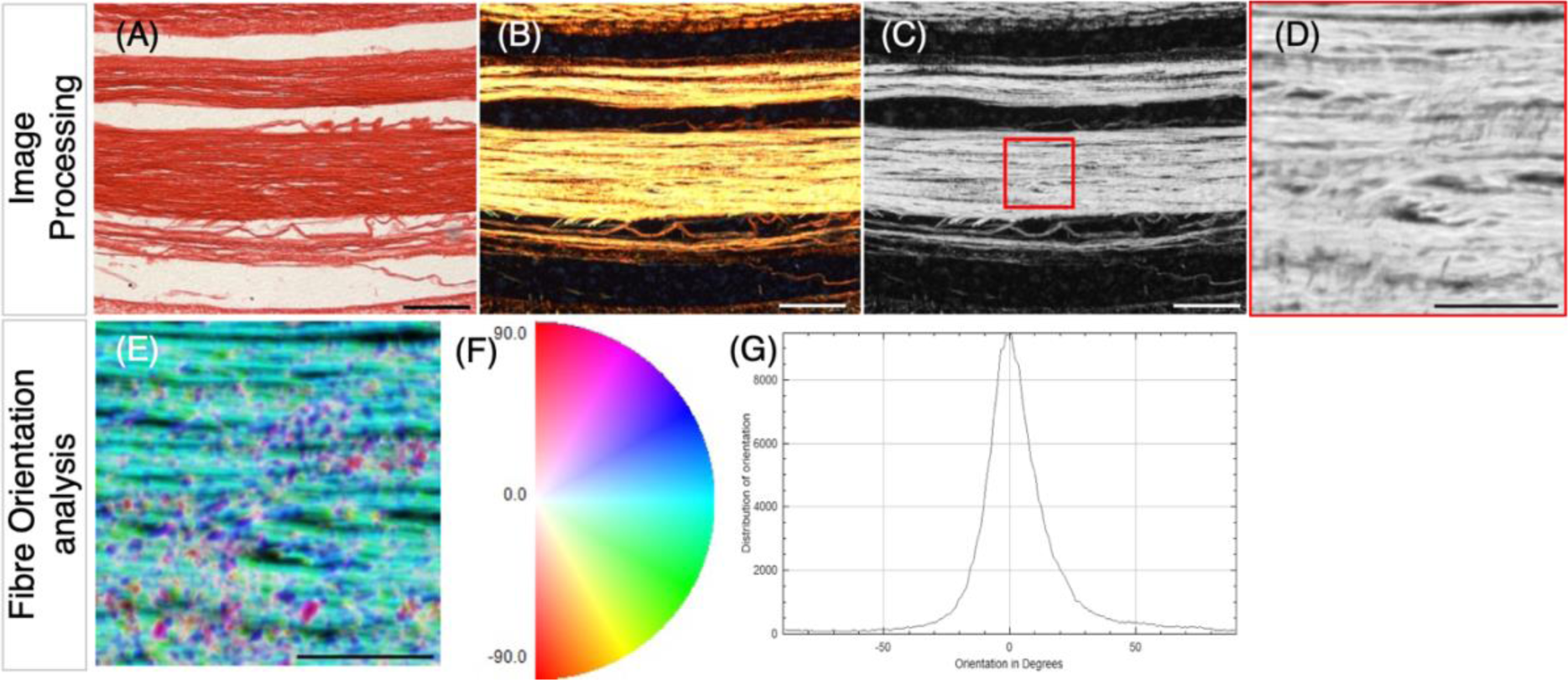
Methodology for collagen fibre orientation analysis following picrosirius red staining of tendon sections and Polarised Light Microscopy (PLM). A, Example brightfield image of longitudinal section through the metatarsal tendons (E18) stained with picrosirius red. B-D, polarised light images of the tendon shown in A. B, shows the merge of two polarised light images; the second taken following rotation of the analyser filter by 45⁰ (to capture more of the birefringence). C, is the 8 bit, greyscale conversion of B showing the selection of the 500 x 500 µm Region of Interest (ROI) for analysis, placed within a fibre bundle (red square). D, shows the selected ROI. E-G, Analysis of fibre orientation within the ROI shown in D using OrientationJ plugin with ImageJ. This generates a fibre orientation colour map (E), where blue/cyan indicates fibres oriented close to 0⁰ and pink/red indicates fibres oriented ∼90⁰ (F). G, Using the colour map information, a histogram of local angles can be generated representing the mean and standard deviation of orientations for that ROI. Data was extracted from the generated tables to represent the proportion of fibres in orientation groupings (within +/-10⁰; +/-10-20⁰ ; +/-20-30⁰) as shown in box plots in figures 2 and 4. Scale bars; A-C 500µm, D and E, 200 µm.

### 2.4 Immunohistochemistry and Collagen detection

Heat-mediated antigen retrieval was performed on histological sections to improve detection for all of the following target proteins, using 0.01 M sodium citrate (pH 8) for 20 mins at 90°C. Collagen was detected using a fluorescently labelled CNA35 collagen binding protein (CNA35-eGFP) (plasmid DNA kindly provided by Maarten Merkx). The recombinant protein was expressed and purified from E. coli (1-2 mg/ml) and utilised as previously described(Mohammadkhah et al., 2017). Blocking was performed using 1% bovine serum albumin (BSA) in PBS for 1 hr at RT, incubation in binding protein was carried out at a dilution of 1:20 - 1:50 in blocking solution, overnight at 4⁰C. Following subsequent PBS washes mounting with DAPI nuclear stain was performed (Prolong Gold antifade reagent with DAPI, Life Technologies). Following microscopic imaging (detailed below), fluorescent intensity analysis was carried out with ImageJ using x20 magnification with a minimum of 5 ROI per image, and 5 technical replicates per biological sample; the mean gray-level representing fluorescence intensity. For YAP and pYAP tissue level detection, following antigen retrieval, paraffin sections were processed, undergoing heat-mediated antigen retrieval, as previous, blocked in 5% goat serum in TBS with 0.1% Tween (TBST) and primary antibody incubated overnight at 4°C. Primary antibodies used were: mouse monoclonal anti-YAP ((63.7) Santa Cruz-sc101199, 1:200) and rabbit polyclonal anti-phospo-YAP, Ser127 (Cell signalling, cat. No 4911, 1:200). Colorimetric secondary detection was performed for whole tissue level analysis using goat anti-mouse (Cell signalling-7056S; 1:2000) or anti-rabbit IgG-Alkaline Phosphatase (Invitrogen-G21079; 1:200 blocking solution) with sections equilibrated to pH 9.5 in NMT buffer (0.1M Tris pH 9.5, 0.05M MgCl_2_, 0.1% Tween or 0.1M NaCl), then developed with NBT (nitro-blue tetrazolium) /BCIP (5-bromo-4-chloro-3-indolyphosphate) diluted in NMT. Colorimetric samples were mounted with Aqua-polymount (Polysciences).

For GAPDH detection, paraffin sections were blocked in 5% bovine serum albumin in PBS, incubated in primary antibody, mouse monoclonal GAPDH (Mouse, Novus Bio, NB300-221 1:200) overnight at 4°C. Fluorescent secondary detection was performed using Alexa Fluor 488 goat anti-mouse IgG (Invitrogen, A11001, 1:250), and mounted with ProLong Gold Antifade Reagent with DAPI (Life Technologies).

Tissue sections were photographed under bright field and fluorescent microscopy (as appropriate) using an Olympus DP72 camera and CellSens software (v1.6).

### 2.5 Image analysis and Quantification of morphological and cellular parameters

The cross-sectional areas of the Achilles tendon, flexor digitorum longus (FDL) and the flexor digitorum brevis (FDB) digit II tendons were quantified from embryos at E20 (HH45) following rigid and flaccid immobilisation and experimental controls (n=7-12 control, n=8-11 rigid and n=7-8 flaccid immobilised biological replicates from 2-4 independent experiments) using measurements from 7-20 adjacent Masson Trichrome stained cross-sections through a 560-1,200 µm portion of the medial tarsometatarsal region of interest (ROI) or each replicate embryo. The boundary of collagen positive (blue) staining defined the boundary of the tendon for measurements (as shown in Figure 3A). The cross-sectional area of the Achilles tendon, FDL2 and FDB2 tendons were quantified independently using ImageJ (v2.1.0/1.53c). The width of the Achilles tendon was also measured (Supplementary Figure 1). The same medial ROI was used to assess the spacing between the FDL and FDB tendons following immobilisation. For each section from each sample 3-8 measurements were taken across the extend of the interface (where approximately parallel) between the tendons, and averaged, to represent the spacing between tendons. Spacing measurements represent n=7 control, n=8 rigid, and n=7 flaccid biological replicates. Effects of immobilisation were also analysed in the knee region, specifically on the patellar tendon (Supplementary Figure 1)

Nuclear imaging and analysis was performed using x 100 magnification images of DAPI stained tendons across normal development E14-E20 and following immobilisation using ImageJ. Cell densities were quantified using the cell counter function in a field of view equal to 137.412µm^2^ (Figure 4C) across 3-5 technical replicates per developmental stage (embryos analysed per stage: E14 n=4, E16, n=3, E18, n=4, E20 n=3) or treatment group (embryos analysed per group; Control n=12, rigid, n=7, flaccid, n=6). Analysis of nuclear shape was performed with original files (Supplementary Figure 3a) cropped to define an ROI of 5µm x 4µm (Supplementary Figure 3b). Cropped images were split into component colour ‘channels’, the blue channel (Supplementary Figure 3c) was determined to have the best signal-to-noise ratio and so was used for further analysis. Images were converted to 8-bit by setting a minimum threshold, using the auto threshold function as a guideline. This converted every pixel above the threshold in the greyscale blue channel to black, and every pixel below the threshold to white (Supplementary Figure 3d). The appropriate threshold values were determined independently for each biological replicate, due to the unavoidable variability in staining contrast or intensity, and in image capture. Any background noise, or out of focus nuclei were removed either manually or using the despeckle function, and converged nuclei were separated (Supplementary Figure 3e). Nuclei were measured using the Analyse particles function (Supplementary Figure 3f), with a minimum particle size of 0.05μm^2^. Nuclei touching the edge of the ROI were excluded to prevent inaccurate shape measurements. The analyse particles function generated shape measures of circularity and aspect ratio. Circularity was calculated as 4π times the area divided by the perimeter squared. A perfect circle has a circularity value of 1.0; as the shape becomes more ellipsoid, its circularity value decreases (Supplementary Figure 3g).

The transverse distance between cell nuclei in adjacent aligned columns in the longitudinal tissue plane were calculated from 5µm x 4µm cropped images for each developmental stage and treatment group. A minimum of 3 images per biological replicates were used to assess the distance between nuclei. Measurements were made between the edges of adjacent nuclei using the line and measure function in ImageJ and each individual measurement was treated independently. The total number of measurements across independent biological replicates were as follows: E14 =110 from 4 biological replicates, E16 = 95 from 3 replicates, E18 = 106 from 3 replicates, E20 = 87 from 3 replicates, Control = 260 from 12 replicates, Rigid immobilised = 156 from 7 replicates and flaccid immobilised = 104 measurements from 6 replicates.

### 2.6 Protein and mRNA Quantification from dissected tendons

Tendons ventral to the ankle joint, including the flexor halluces brevis, the flexor perforan et perforates and the flexor digitorum were sub dissected from hindlimbs between the ankle joint and proximal to the footplate. Tendons were pooled from each embryo and placed in TRIzol (Thermo Fisher Scientific), stored at -70°C. Mechanical homogenisation was performed using polypropylene pestles and high speed vortexing. Protein extraction was performed according to the manufacturers protocol (TRIzol; Thermo Fisher Scientific) and resuspended in 8M Urea with 1% SDS in 0.05M Tris (pH=8) solution, quantified (Qubit 2.0 fluorometer, Invitrogen) and stored at -20°C. For western blotting, 40 µg of protein were separated on a 10% pre-cast polyacrylamide gel (Bio-Rad) and transferred to PVDF membranes. PVDF membranes (0.45 µm, Bio-Rad Transblot Turbo PVDF (Cat#10026934)) were blocked with 5% milk powder in TBST and incubated in 1:1000 anti-YAP ((63.7) Santa Cruz-sc101199), 1:1000 anti-phospo-YAP (cell signalling, cat. No 4911) or 1:1000 anti GAPDH (NB300-221 Novus Bio). Detection and visualisation was performed using fluorescent detection with Li-Cor antibodies (1:10,000 goat-anti-mouse IRDye 680RD and goat-anti-rabbit IRDye 800CW) and a Li-Core Odyssey system (Odyssey XF). Densitometry analysis was performed using ImageJ software for YAP and pYAP protein levels normalised to GAPDH protein levels, comparing developmental timepoints (E14-E20) and between control and immobilised samples at E20 (n=12 (control), n=11 (rigid), n=3 (flaccid) independent embryos in each case).

RNA was extracted from the stored TRIzol tissue homogenate with chloroform and purified with a TRIzol Purelink kit (Invitrogen) following manufacturers’ instructions. Typically, mRNA was re-suspended in 30 µl of nuclease-free water and quantified using the Qubit 2.0 Quantitation System (Invitrogen). mRNA was reverse transcribed using a standard quantity of total RNA (500 ng) and High Capacity cDNA Reverse Transcription Kit (Applied Biosystems™: 4368814), diluted 1:5 with RNase free water. Chick primers for genes of interest (Table 1) were synthesised by Merck. *Gapdh* was used as the normaliser transcript. Real-time PCR quantification was performed using an ABI 7500 Sequence Detection system (Applied Biosystems) using SYBR green gene expression quantification (Applied biosystems) 5 μl of cDNA preparation was diluted 1:5 with RNase free water, 10 μl of 2x SYBR green PCR master mix (Applied Biosystem), 0.5 µl (10 μM) of each primer and 4 µl RNase free water). Samples were assayed in triplicate in one run (40 cycles), which was composed of three stages, 95°C for 10 min, 95°C for 15 s for each cycle (denaturation) and 60°C for 1 min (annealing and extension). Data was analysed using relative quantification and the Ct method as described previously (Livak and Schmittgen, 2001). The level of gene expression was calculated by subtracting the averaged Ct values (Ct is the threshold cycle) for *Gapdh* from those of the gene of interest. The relative expression was calculated as the difference (ΔΔCt) between the Ct of the test stage or condition (immobilised) and that of the compared stage or the control condition. The relative expression of genes of interest were calculated and expressed as 2-ΔΔCt. Relative quantification values are presented as fold changes plus/minus the standard error of the mean relative to the comparator group, which was normalised to one.

**Table 1:**
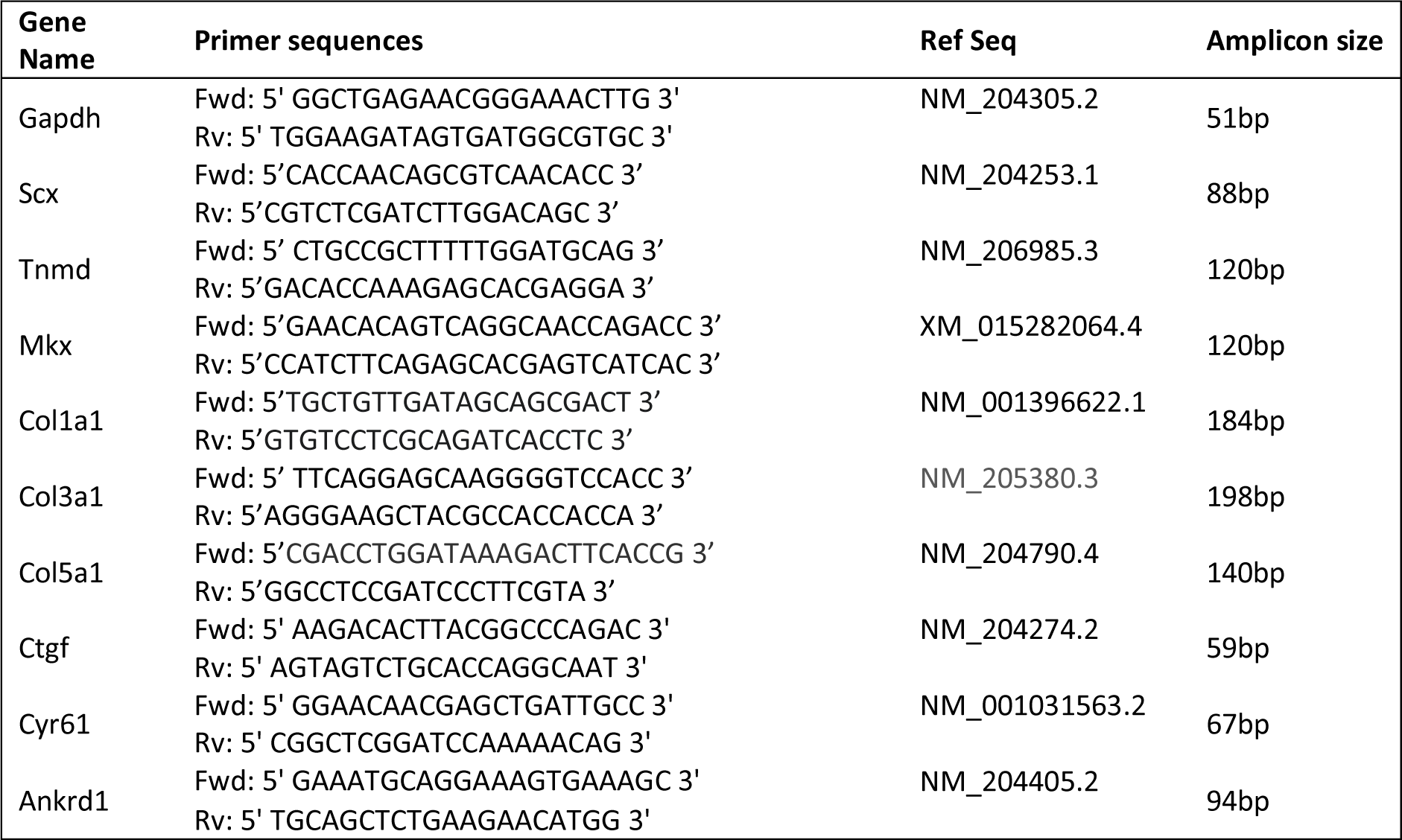
Details of primers used in qRT-PCR.

### 2.7 Statistical analysis

A range of 3 -6 independent biological replicates (embryos) were analysed for all normal development stages and between 8 - 15 independent biological replicates for control and immobilised specimens across 2 - 4 independent experiments. Specifically for morphological analysis 12 controls, 11 rigidly and 8 flaccidly immobilised biological replicates were assessed. For PLM 14 controls, 7 rigidly and 6 flaccidly immobilised biological replicates were measured. For nuclear comparisons 12 control, 7 rigidly and 6 flaccidly immobilised biological replicates were assessed, with 604 nuclei from control, 356 nuclei from rigid and 308 nuclei from flaccid immobilised replicates analysed for nuclear shape. For all data SPSS statistics (IBM^®^, v.27) was used for statistical analysis. Significance was determined by one-way ANOVA, followed by Tukey’s post hoc test with a 95% confidence interval. P values of ≤ 0.05 were considered significant.

## 3 Results

### 3.1 Collagen fibre alignment increases during tendon maturation, as tenocytes stack into parallel columns

From the earliest stage examined (E14), abundant collagen deposition was detected on longitudinal sections through metatarsal tendons (Figure 2A), with no apparent difference in collagen level across subsequent ages (E14-E20) revealed by fluorescent intensity analysis (data not shown). To look closer at deposited collagen over time, picrosirius red staining (PSR) combined with polarised light microscopy (PLM) was used to analyse the spatial organisation of the collagen network(Liu et al., 2021), in particular permitting quantitative analysis of collagen fibre alignment as well as reflecting the maturity of the deposited collagen(Patel et al., 2018); the colour of the birefringent light has been previously taken to indicate increasing fibre thickness and maturity from green to yellow, orange and red. Figure 2B shows representative images of PSR-PLM of metatarsal tendons across stages of development indicating that collagen fibres shifted from a combination of green and yellow birefringence at E14 and E16 to predominantly yellow at E18 and E20. Some red birefringence is visible at the edges of tendons from E18 (Figure 2B). Analysis of the alignment of fibres shows progressively more alignment across late development (Figure 2C). Figure 2C represents the proportion of fibres in orientation groups; +/-10⁰, +/-10⁰-20⁰ and +/-20⁰-30⁰ of the longitudinal axis across time and shows that the proportion of fibres in the +/-10° category increases significantly (e.g. E14-E20: 41.7 ± 2.2% to 59.8 ± 2.1% p<0.001) with an accompanying reduction in the proportion of fibres in the third category, ±20° -30° (e.g. E14-E20 13.3±1% to 7.7 ±0.6% p<0.001). This shift is also demonstrated by the proportion of fibres represented within +/-30⁰ of the longitudinal axis, increasing from 80% at E14, 83% at E16, 88% at E18 to 91.5% at E20.

The number of cells per unit area within the tendons reduced from E14 to E18 (p=0.044), E14 to E20 (p=0.005) and from E16 to E20 (p=0.024) (Figure 2E), with the outline shape of the nuclei becoming more circular (value closer to 1) across development from E14 to E20 (Figure 2F), illustrated by DAPI staining and nuclear aspect ratio (Figure 2D). At E14 the greatest diversity of nuclear shapes was identified, with only 20.1% of nuclei with an aspect ratio of <2 (i.e. more circular; lightest blue in Figure 2G). As development proceeds the diversity in nuclear shape reduces, with 91.4% of nuclei in the most circular outline category at E20 (Figure 1G). In addition to the more uniform nuclear shape, the cells have aligned into more organised columns with the distance between nuclei between separate longitudinal columns increasing significantly between each pair of consecutive stages examined (Figure 2H). The mean distance between nuclei increased from 0.1 ± 0.005μm at E14, to 0.38 ± 0.02μm at E20 (p≤0.001, Figure 2H). In summary, our data show that in embryonic chick ankle tendons collagen fibres become more aligned, cell density reduces, the distance between columns of cells increases and nuclei become more uniformly circular as they mature across late development.

**Figure 2:**
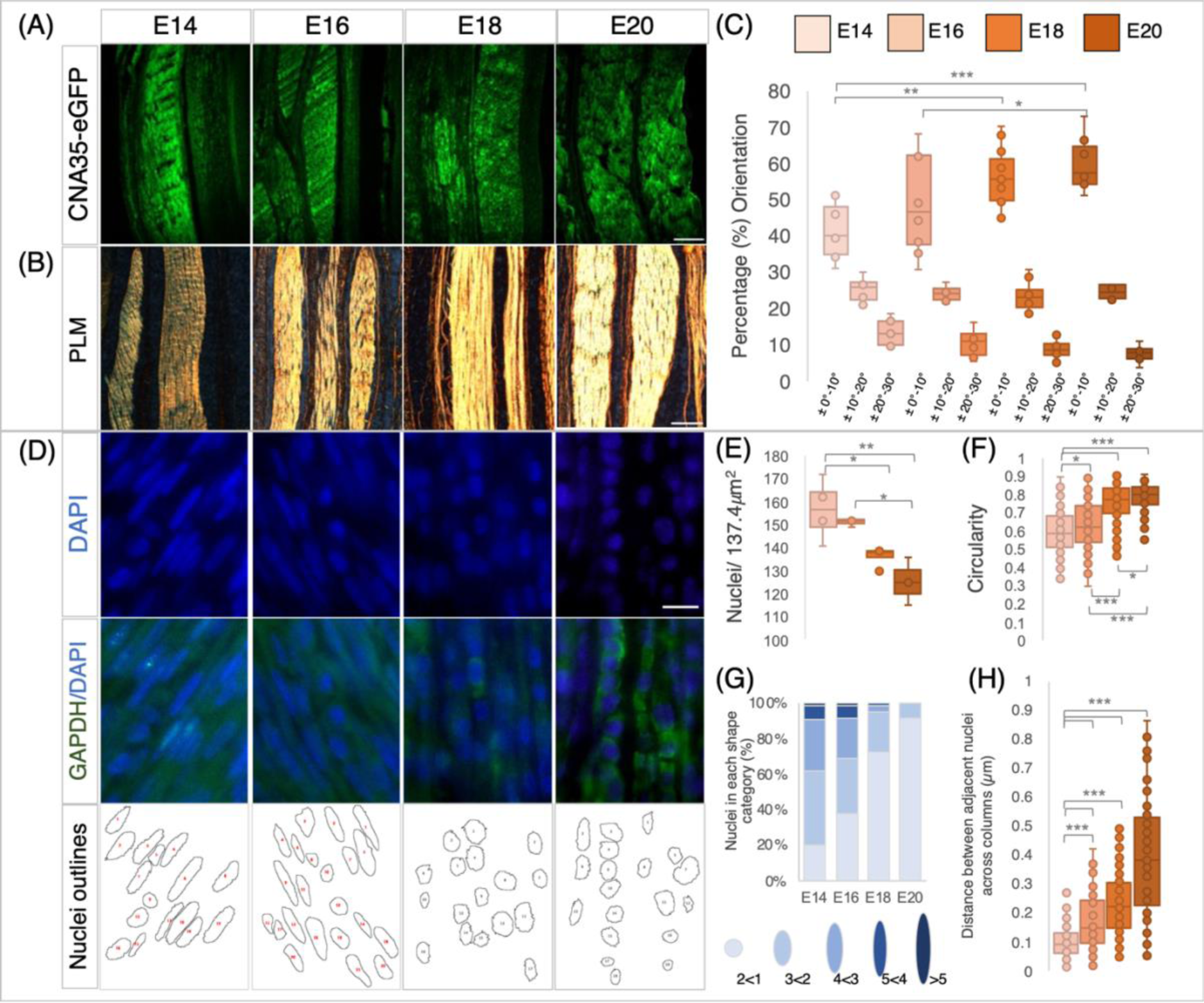
Changes in collagen deposition and cellular organisation in chick metatarsal tendons across late stages of development. A, Visualisation of collagen deposition in longitudinal sections through embryonic chick metatarsal tendons using a fluorescently tagged pan collagen binding protein (CNA35-eGFP) at developmental time points indicated. B, Collagen fibre bundles visualised by picrosirus red staining and polarised light microscopy (PLM) in longitudinal sections of metatarsal tendons. C, Collagen fibres become more aligned across development: Quantification of fibre orientation was carried out using the Orientation J plugin of Image J to analyse birefringence patterns, as illustrated in Figure 5. The box plot represents the proportion of fibres within three orientation groupings with respect to the tendon long axis: +/-0° -10°, +/-10° - 20°, +/- 20° - 30° (measurements taken from n=4 (E14), n=3 (E16), n=5 (E18) and n=4 (E20)). D, Representative images of sections across the developmental profile showing nuclei (DAPI), cytoplasm (GAPDH) and images merged (GAPDH/DAPI). Outline drawings of nuclei extracted from these images are shown in the bottom row. E, Box plot representing cell density per 137.4μm^2^ regions quantified from multiple sections through the central body metatarsal tendons across development (n=4 (E14), n=3 (E16), n=4 (E18) and n=3(E20) embryos). F, Nuclear circularity increases (value closer to 1) during tendon maturation, (n=219 (E14), n=174 (E16), n=190 (E18) and n=151(E20) cell measurements across replicates per stage). G, Nuclear shape becomes more uniformly round during tendon maturation: Histogram representing the proportion of nuclear shapes (schematic representations with aspect ratio values shown below for each category) found in embryonic tendons. H, The distance between cell nuclei in adjacent aligned columns increases significantly across development (n=110 (E14), n=96 (E16), n=106 (E18) and n=87 (E20) measurements across replicates/ stage). Plot whiskers represent min and max data values in all box plots. *\*p≤0.05, **p≤0.01, ***p≤0.001.* Scale bar in (A) 200μm, (B) 500μm, (D) 1 μm.

### 3.2 Both rigid and flaccid paralysis during tendon maturation led to reduction in size of multiple tendons, while rigid paralysis also reduced tendon spacing

We previously showed that rigid paralysis induced by treatment with Decamethonium bromide (DMB) from E14 to E17 caused a reduction in the size of metatarsal tendons(Peterson et al., 2021). Here we examine both rigid and flaccid (induced by treatment with Pancuronium Bromide (PB)) paralysis across an extended period during which the tendons are maturing (E16-E20), also analysing effects on multiple tendons. Both rigid and flaccid paralysis significantly reduced the size of the Achilles, Flexor Digitorum Brevis (FDB) and Flexor Digitorum Longus (FDL) digit two tendons at E20 (HH45), in the tarsometatarsal region (Figure 3). Significant decreases in tendon size were observed through measurements of cross-sectional area of the Achilles and both Flexor digitorum tendons (Figure 3D, G). Effects on the Achilles tendon were also assessed through width measurements at defined points, confirming the finding of significant decrease in size following both types of paralysis (Supplementary Figure 1). We also examined the effect of paralysis at the knee joint region and show reduction in the width of the patellar tendon (ligament) (Supplementary Figure 1).

Paralysis also affected the spacing between the FDB and FDL tendons but here rigid and flaccid paralysis had different effects. While there was a severe effect on spacing following rigid paralysis (Figure 3F; p<0.0001, n= 8 immobilised embryos; n= 7 control, representative section shown in Figure 3B, green arrow), no significant reduction was observed following flaccid paralysis (n= 7 each of flaccid and control embryos). However, there was a larger variation in inter-tendon spacing measurements with flaccid paralysis (average of 23.9 ± 2.7 μm compared to 34.3 ± 0.7 μm in controls).

**Figure 3:**
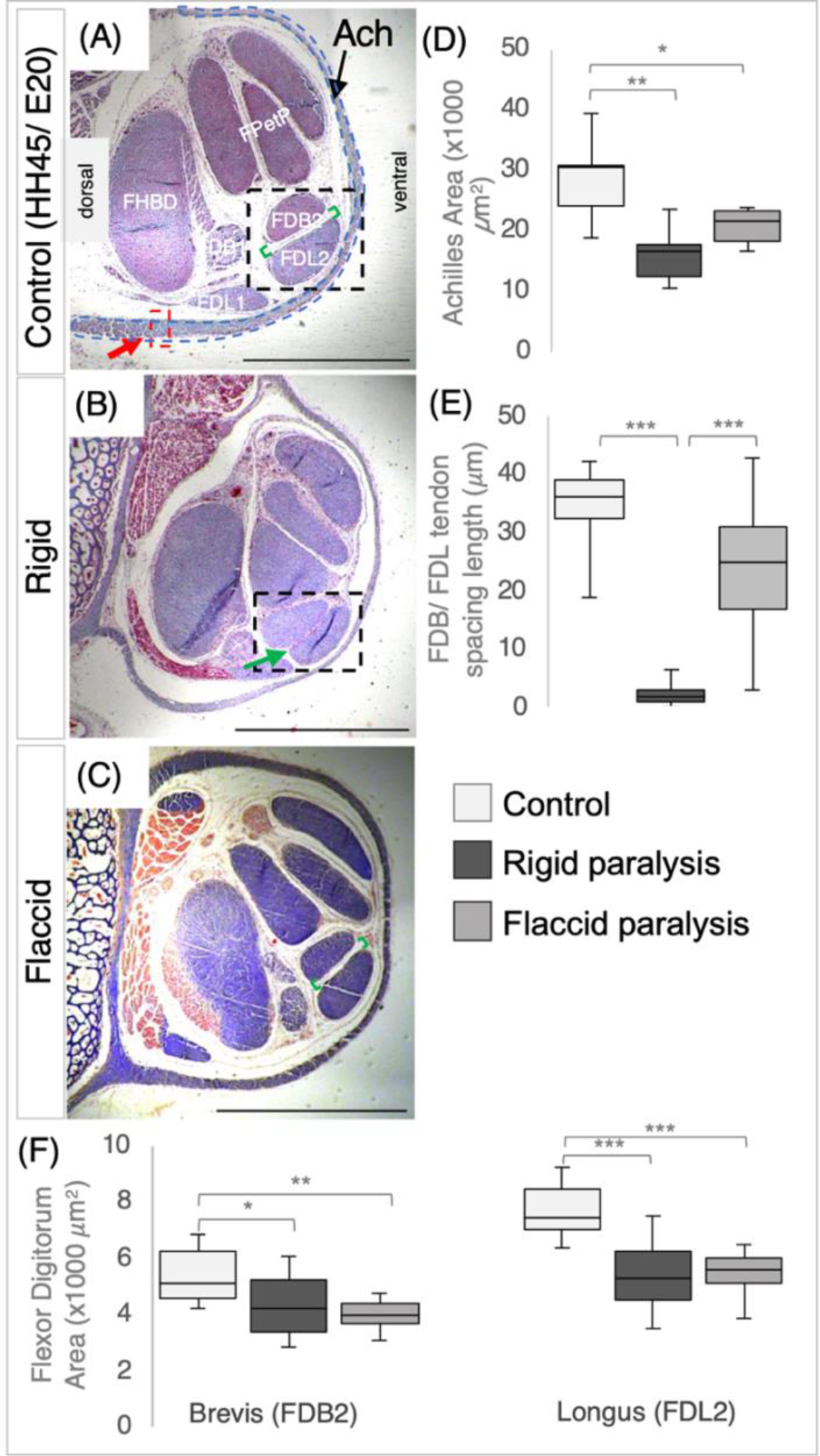
Immobilisation by both rigid and flaccid paralysis alters tendon size while only rigid paralysis alters spacing between adjacent tendons. A-C, Representative cross sections of control (A), rigid (B) and flaccid immobilised (C) tarsometatarsal regions of chick hindlimbs at embryonic day 20 (E20)/ Hamburger and Hamilton stage 45 (HH45) stained with Masson Trichrome. D, Box plot of cross-sectional area of Achilles tendon, with area defined by the dashed blue line indicated in (A). Both area and width were significantly reduced following immobilisation (n=7 (control), n=6 (rigid), n=6 (flaccid)). E, Box plot representing the inter tendon spacing (shown with green brackets in A and C) across groups, a significant reduction in spacing following rigid immobilisation is shown by the green arrow in B, (n=7 (control), n=8 (rigid), n=7 (flaccid)). F, Box plots of cross sectional area measurements of the FDB2 and FDL2 showing significant reductions following immobilisation (n=12 (control), n=11 (rigid), n=8 (flaccid)). Plot whiskers represent min and max data values in all box plots. *\*p≤0.05, **p≤0.01, ***p≤0.001.* Scale bars (A-C) 1000μm, Achilles (Ach), Flexor Digitorium Longus/Brevis Digit I (FDL/B1), Flexor Digitorium Longus/Brevis Digit II (FDL/B2), Flexor perforan et perforates Digit III & IV (FPetP), Flexor Halluces Brevis Digits II, III, IV (FHBD).

### 3.3 Rigid and flaccid paralysis have differential effects on collagen fibre alignment and cellular organisation

While there was no visual effect on deposited collagen following immobilisation (Figure 4A) collagen fibre alignment within metatarsal tendons was differentially affected by rigid and flaccid paralysis from E16-E20. The intensity of fluorescent detection across conditions was not altered and the birefringence pattern shows similar colours, with the majority of the fibres appearing yellow (with no discernible green) indicating intermediate maturity, and with red (indicative of more mature) appearing at the margins of all tendons. Following fibre alignment analysis, the box plot in Figure 4B groups the proportion of fibres in orientation categories with respect to the long axis of the tendon. Rigid immobilisation resulted in a higher proportion of fibres within +/- 10⁰ of the long axis (66.9 ± 1.5% compared to 56.6 ±1% in the control group; p<0.001) while flaccid paralysis had the opposite effect with a lower proportion aligned (50.7 ±2%, p=0.011) (Figure 4B). The same patterns of change were observed in all categories.

Examining the effects of immobilisation on tendon cellular organisation, both types of immobilisation had effects but to different extents, more pronounced with rigid paralysis. One distinct difference was observed in cell density analysis with rigid paralysis leading to an increase in the number of nuclei per unit area compared to both normal moving controls (p=0.007) and flaccidly (p=0.027) immobilised tendons (Figure 4C, G). Flaccid paralysis had no significant effect. Rigid and flaccid paralysis were both found to alter nuclear shape, with nuclei becoming more elongated compared to controls at E20/HH45 (p<0.001 in both immobilisation modes), however, even though nuclei under flaccid conditions are elongated, they are less so than rigid nuclei (Figure 4D, H, p=0.001). While nuclei are arranged in columnar parallel patterns across treatment groups, the shape of cells following immobilisation was disturbed, as revealed by cytoplasmic staining of GAPDH (Figure 4E). In control tendons, the tenocytes are regularly stacked and clearly defined with a cuboidal shape and clear separation between neighbouring cells (Figure 4F orange arrowheads) whereas following immobilisation there is no visual separation between cells within longitudinal columns. Additionally, the transverse distance between nuclei in adjacent aligned columns is reduced under both immobilisation conditions compared to control (rigid 0.16 ± 0.01μm, flaccid 0.22 ± 0.01μm compared to control 0.36 ± 0.01μm ((p≤0.001, Figure 4I), with rigid paralysed embryos being more severely affected.

In line with the observed loss of regular cellular organisation, the diversity of nuclei shapes significantly increases following immobilisation [Supplementary Figure 2]. Disruption of collagen fibre alignment and cellular organisation with immobilisation, strongly indicates that structural organisation of tendons is reliant on movement.

**Figure 4:**
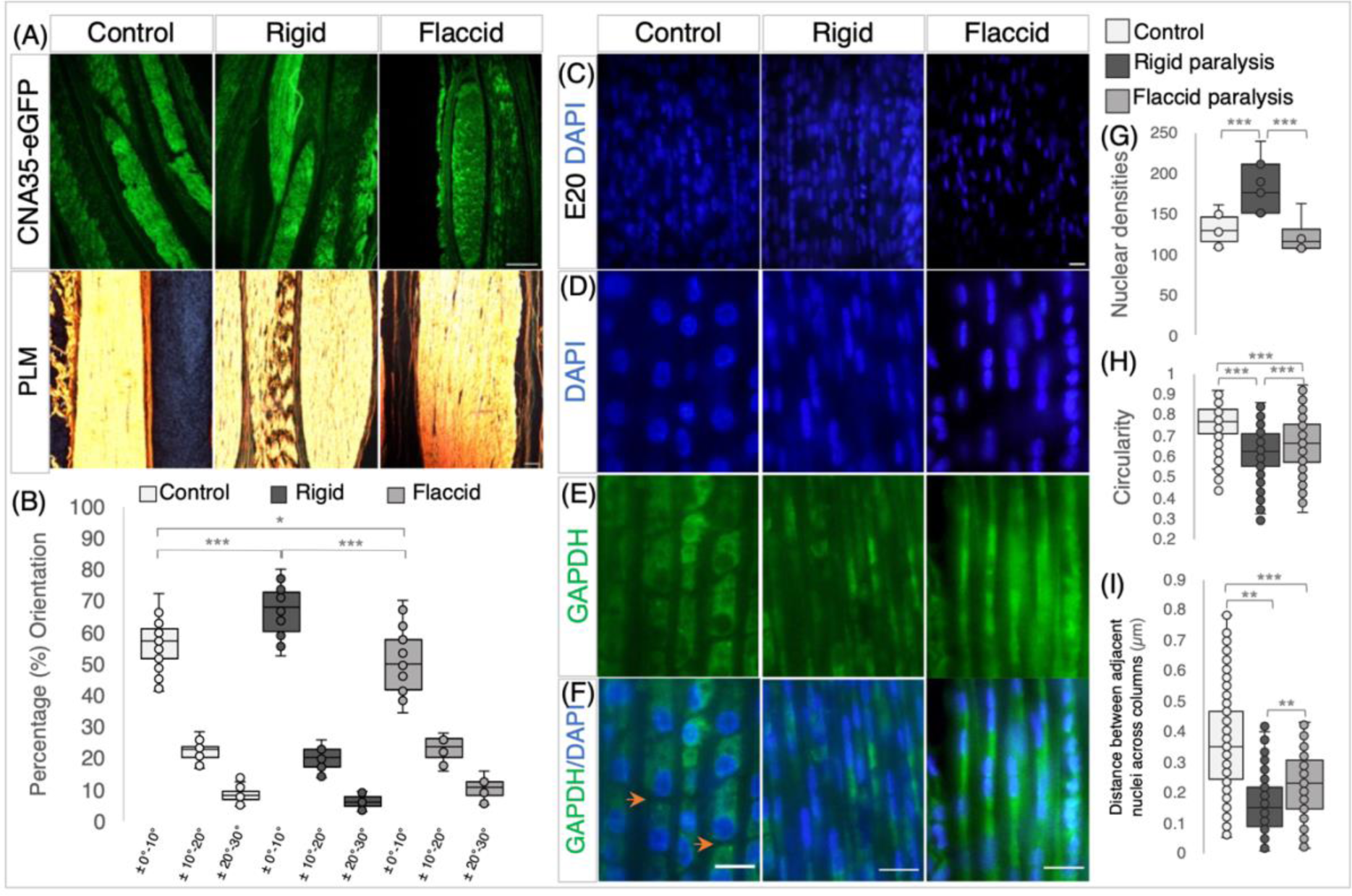
Rigid and Flacid paralysis have differential effects on **c**ollagen fibre alignment and cellular organisation. A, Collagen visualisation using collagen binding protein (CNA35-eGFP), with polarised light microscopy (PLM) following picrosirus red staining (below) on longitudinal sections of metatarsal tendons at E20/HH45, following rigid and flaccid paralysis compared to control embryos, as indicated. B, Quantification of collagen fibre orientation shows that fibre alignment is altered by immobilisation, with increased alignment following rigid paralysis and less alignment with flaccid paralysis. The box plot represents the proportion of fibres within three orientation groupings with respect to the tendon long axis: +/- 0° - 10°, +/-10° - 20°, +/- 20° - 30° (measurements taken from n=14 control, n=7 rigid, n=5 flaccid immobilised embryos). C, Representative views of DAPI stained nuclei in E20 metatarsal tendons from control, rigid and flaccid immobilisations within the 137.4μm^2^ field of view used for cell counts. D-F, higher magnification views demonstrating nuclear shape D, GAPDH immunodetection (cytoplasm) (E) and merged images (F) across treatment groups. Orange arrowheads highlight separation between cells in the longitudinal axis (F). G, cell density increased with rigid paralysis (n=12 control, n=7 rigid, n=6 flaccid immobilised embryos) while flaccid paralysis had no detectable effect. H, Nuclear shape is less circular following immobilisation compared to controls, with a more pronounced effect following rigid than flaccid paralysis, (dots indicate cell replicates: n=604 control nuclei from n=12 embryos, n=356 rigid nuclei from n=7 embryos), n=308 flaccid nuclei from n=6 embryos) (see D). I, The distance between cell nuclei in adjacent aligned columns of cells within the tendon decreased significantly following immobilisation with a greater reduction following rigid paralysis (dots indicate individual measurements; n=260 (control), n=156 (rigid), and n=106 (flaccid). Plot whiskers represent min and max data values in all box plots. *\*p≤0.05, **p≤0.01, ***p≤0.001.* Scale bar 200 μm (A and B), 1 μm (C - F).

### 3.4 Components of the YAP signalling pathway are expressed in the late maturing tendon with alteration of target gene expression levels following immobilisation

To investigate the YAP signalling pathway as a possible mechanoregulator of tendon maturation we set out to establish if YAP pathway components are localised in the tendon during late development and if levels are altered following immobilisation, comparing rigid and flaccid paralysis. Using immunodetection on tissue sections we could readily detect both the active form (YAP) and inactive form (pYAP) in the metatarsal tendons across time points and following immobilisation (Figure 5A; n=8 rigid, n=6 flaccid biological replicates histologically analysed). No consistent differences in localisation patterns were observed; any apparent differences between adjacent tendons are likely due to the precise plane of section and were not consistently reproduced (n= 12). This was supported by quantification of YAP and pYAP absolute protein level accumulation via western blot which showed no significant change in levels of either protein, but it revealed a trend of reduction in mean YAP levels across later stages of development, between E14 - E18 and E16 - E18 (Figure 5B, #p=0.067 and p=0.059 respectively). To check for a possible shift in the relative levels of YAP and pYAP the ratios of active YAP normalised by total YAP (YAP+pYAP) levels were examined and showed no change over time (YAP/(YAP+pYAP) ratios: E14, 0.44 ± 0.06; E16, 0.52 ±0.007; E18, 0.38 ± 0.05; E20 0.53, ±0.16). Similarly, no quantitative differences were detected for YAP or pYAP protein levels following immobilisation (Figure 5C, histogram representative of n=15 controls, n=11, rigid n=3 flaccid embryos screened for YAP, YAP (YAP+pYAP) =0.51 ± 0.04 for control and 0.48 ± 0.04 for immobilized).

Gene expression profiling of in vivo tarsometatarsal chick tendons every 24 hours from E14 to E20 show that the relative expression of *Scx* increases between E14 and E16 (p=0.007) and then reduces from E16 to E17-E20 inclusive (Figure 5D, (E16-E17 p=0.01, E16-E18 p=0.01, E16-E19 p=0.026, E16-E20 p=0.001)). Similar decreases in *Tnmd* expression were observed between E16 to E18 and E20 (p=0.02 and p=0.01, respectively) and E17 to E18 and E20 (p=0.02 and p=0.01 respectively, Figure 5D). No significant changes in gene expression were observed for *Mkx*, or tendon specific collagen genes (*col1a1*, *col3a1* or *col5a1*) across the late stage of tendon development. Interestingly, YAP target genes were found to be expressed across developmental time, from E14 to E20, indicating pathway activity, with *Cyr61* and *Ankrd1* more highly expressed at later stages (Figure 5D). An increase in *Cyr61* was observed between E14-E17 (p=0.044), E14-E20 (p=0.002) and E16-E20 (p=0.022) and increases in expression from E14-E19 and E16-E19 for *Ankrd1* (p=0.042 and p=0.03 respectively) (Figure 5D). Detection of pathway components and quantitative changes in expression levels indicate that the pathway may be modulated in this developing system.

Multiple disturbances in gene expression levels were detected with immobilisation (Figure 5E). Rigid immobilisation resulted in more gene disturbances than flaccid immobilisation compared to control expression (5 and 2 genes, respectively), with four of the rigid disturbances identified also being different to flaccid expression (*Tnmd*, *Mkx*, *Ctgf* and *Cyr61*); suggesting that rigid and flaccid immobilisation may affect the YAP pathway differently (Figure 5E). Alteration in the expression of tendon specific genes *Scx*, *Tnmd* and *Mkx* were observed; a 2.21 fold reduction in *Scx* with flaccid paralysis (p=0.006), while a 2.29 fold and 2.68 fold increase in expression of *Tnmd* and *Mkx* were identified with rigid paralysis (p=0.01 and p=0.012 respectively, Figure 5E). No expression changes were observed in any of the collagen genes screened. Rigid immobilisation led to increased expression of YAP target genes *Ctgf* (p=0.000002), *Cyr61/Ccn1* (p=0.000117) and *Ankrd1* (p=0.000263). Flaccid immobilisation led to an increased expression of *Ankrd1* only (p=0.001), while both *Ctgf* and *Cyr61* expression levels were not different to control levels but significantly less than following rigid immobilisation (Figure 5E). The observed increased activity of the pathway in rigidly immobilised tendons suggests that YAP pathway output is tightly controlled during normal development, mediated by mechanical cues and/or biophysical changes, impacted by immobilisation; most strongly when constant static muscle forces are imparted but dynamic stimulation is lost (rigid paralysis) compared to flaccid paralysis in which both the dynamic and static muscle forces are removed.

**Figure 5:**
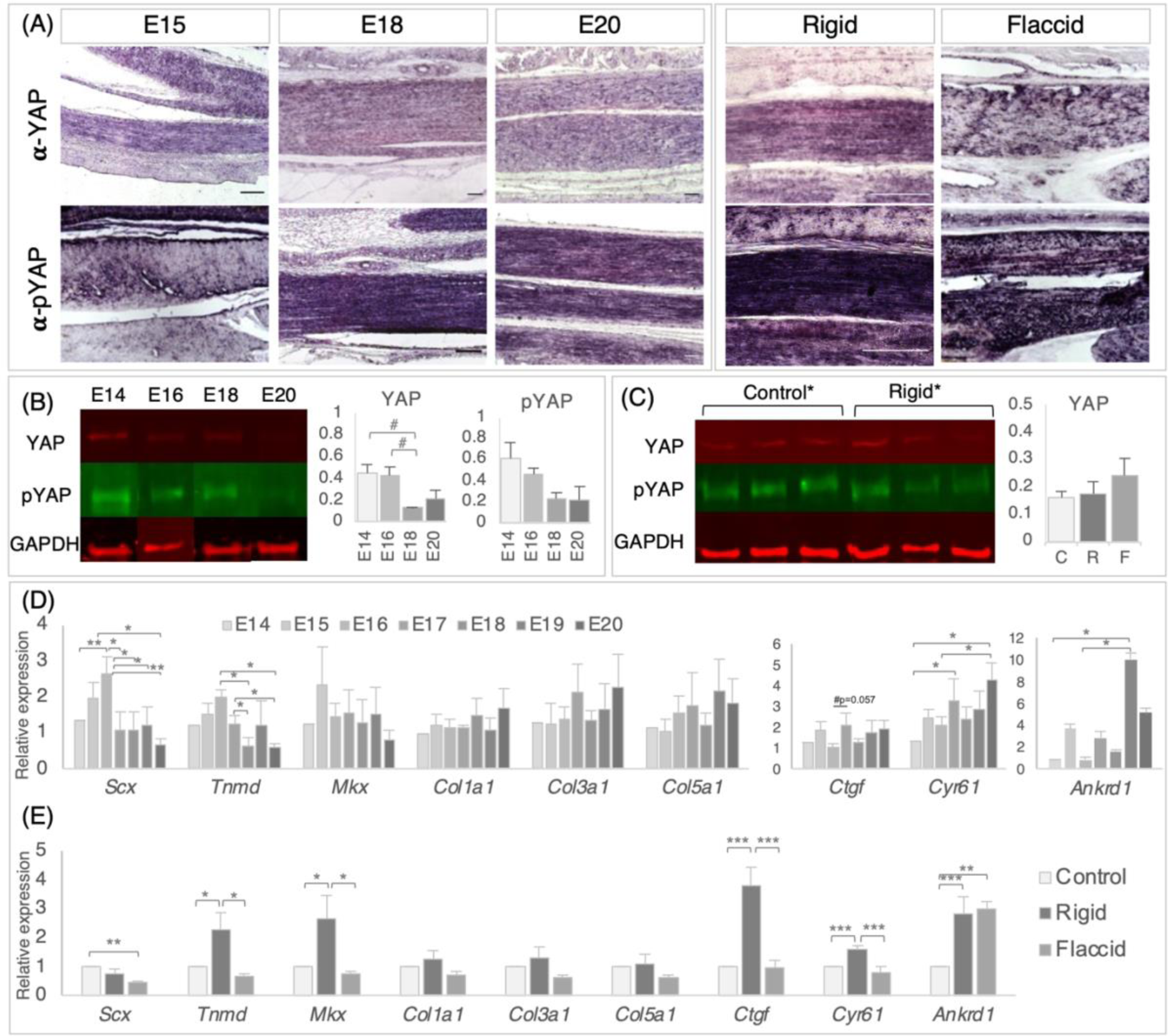
Detection and quantification of YAP signalling pathway components in the late stage tendon, across development and following immobilisation, shows that target gene expression is altered following immobilisation. A, Immunodetection of YAP and pYAP proteins show widespread localisation throughout late stage developing metatarsal tendons, across developmental time points and following immobilisation. Example longitudinal sections are shown at E15, E18 and E20 and following rigid and flaccid paralysis at E20. (E20 normal development and E20 control samples were separately analysed and were equivalent throughout, so the E20 column represents both). B, Protein quantification of YAP (65kDa) and pYAP (65kDA) by western blot, normalised to GAPDH (36kDa) levels, over developmental time points, reveals a trend of reduced mean YAP protein levels; between E14-E18 (#p=0.067) and E16-E18 (#p=0.059), but with no significant difference. Example protein bands for each stage (E14, E16, E18 and E20) are shown. The histograms show YAP and pYAP absolute protein levels (n=3 (E14, E18, E20) n=5 (E16)). C, YAP and pYAP quantification following immobilisation show no significant differences in protein levels following immobilisation (3 example replicate protein bands shown for control and rigid groups). The histogram shows data for YAP(n=15 (control (C)), n=11(rigid (R)) n=3 (flaccid (F))). D, Relative gene expression levels in metatarsal tendons across developmental time points as indicated, for *Scleraxis*, *Tenomodulin*, *Mohawk, collagen 1, collagen 3, collagen 5,*together with YAP pathway target genes *Ctgf*, *Cyr61* and *Ankrd1*. All data were normalised to *Gapdh* expression and presented relative to E14 expression levels (n=3-6). Note that YAP target genes *Cyr61* and *Ankrd1* showed significantly increased expression in later stages of development (n=3-6). E, Relative gene expression following rigid (n=11-13) and flaccid immobilisation (n=6). All data were normalised to *Gapdh* levels and expressed relative to control (n=20). While the expression levels of collagen genes were not altered by immobilisation, both tendon regulatory genes (Tnmd, Mkx) and YAP target genes were upregulated by rigid immobilisation but not flaccid. Plot whiskers represent min and max data values in all box plots. ^#^p*≤0.1 *p≤0.05, **p≤0.01, ***p≤0.001.* Scale bars 100μm.

## 4 Discussion

To understand better the tissue level, cellular and molecular changes that take place during tendon maturation we report a two-pronged experimental approach; 1) we profiled changes in vivo across the relevant period of development; E14-E20 in the chick and 2) we immobilised embryos, comparing rigid and flaccid paralysis, to excavate more precisely how maturation is dependent on embryo movement. We show that across the key developmental time points, collagen is deposited with increasingly more mature and aligned fibres; that cell density decreases; and that cell shape becomes increasingly cuboidal with cells stacked end to end into parallel columns within the developing tendon, with increased distance between columns. We show that both rigid and flaccid paralysis affect tendon maturation, but the effects are distinct; while both affect tendon size, only rigid paralysis significantly affects flexor tendon spacing, increases fibre alignment and cell density. Rigid paralysis has a more extreme effect on nuclear shape and spacing between tenocyte columns. To explore the mechanisms involved in a cellular response to mechanical stimulation from movement during tendon maturation we examined components of YAP signalling and show that both YAP (active) and pYAP (inactive) are readily detected in the late developing tendon, although there were no quantitative differences in the levels across developmental time points or under immobilisation. However, YAP target gene expression was upregulated across late development, indicating that the pathway is modulated. Furthermore, following rigid paralysis, expression levels of YAP target genes were further elevated implicating YAP signalling in the mechanoresponse during late tendon development for the first time.

Immobilisation of chick embryos across late stages of tendon development (from E15 or E17) previously showed that mechanical properties of the tendon are affected, with a reduction in the modulus following rigid(Pan et al., 2018; Peterson et al., 2021) and flaccid paralysis(Peterson et al., 2021). Indeed, we previously showed that flaccid paralysis led to a stronger reduction in modulus than rigid paralysis(Peterson et al., 2021). Here we show further distinction between the effects in several respects, with, most notably, rigid paralysis alone causing an increase in fibre alignment and cell densities and a greater effect on YAP target gene expression. In this experimental system rigid paralysis is induced by treatment with DMB, a neuromuscular blocking agent that induces tetanus(Osborne et al., 2002), with sustained contraction of all skeletal muscles, i.e dynamic forces are removed but there are constant static forces(Nowlan et al., 2008), albeit with reduced force given reduced muscle size and contractile properties(Reiser et al., 1988). In the case of flaccid paralysis induced by PB treatment, both dynamic and static forces are absent(Osborne et al., 2002; Rolfe et al., 2017). Therefore, unique or exaggerated effects as a result of rigid paralysis are likely due to the constant strain applied and/or the lack of dynamic stimulation whereas stronger effects of flaccid paralysis indicate features that are more sensitive to absolute reduction in force experienced. For example the exaggerated alignment of collagen fibrils and increase in cell density respond to inappropriate constant mechanical force from muscles in tetanus whereas, the reduction in fibre alignment and increased effect on reduction in tendon modulus by flaccid paralysis(Peterson et al., 2021) indicates a reliance on the level of force applied. This suggests that not only is tendon maturation dependent on the level of muscle force applied, but dynamic stimulation is required for specific aspects. A milder effect of flaccid paralysis compared to rigid paralysis has also been reported previously in the developing chick spine(Rolfe et al., 2017) and in joint formation(Osborne et al., 2002).

Collagen fibres run parallel to the tissue long axis in tendons and ligaments which is a key factor in their high tensile strength and ability to withstand large forces(Birk et al., 1995; Birk et al., 1991). Therefore, the structural organisation of collagen is an important aspect of the system to monitor. Birk et al (1995) showed a large step increase in the length of collagen fibrils by lateral fusion at E17 in the chick(Birk et al., 1995) coincident with increased tensile strength(McBride, 1988), while our previous study showed an increase in the fibril:tissue strain ratio, consistent with an increase in the length of the collagen fibrils(Peterson et al., 2021). Here visualisation of collagen deposition using a fluorescently tagged collagen binding protein showed readily detectable collagen across the developmental profile. Using PLM to assess fibre orientation showed an increasing proportion of fibres within +/-10⁰ of the tendon long axis across time, going from approximately 40% to 60% between E14 and E20 (Figure 2C). Rigid and flaccid immobilisation both affected orientation but in opposite directions; rigid increasing the proportion aligned while flaccid led to a decrease (Figure 4B). This shows that muscle contraction is required for optimum alignment of fibres with the constant static force of rigid paralysis causing increased alignment. This key aspect of collagen structural change, so important for mechanical properties of the tissue, is stimulated by normal embryo movement.

The collagen fibrils are deposited by tenocytes but we know very little about how this later aspect of tenocyte differentiation is regulated, or even how this important maturation process is achieved by the cells, therefore we wished to examine cellular organisation in the maturing tendon in particular. McBride (1985) showed preliminary data (n=2 per stage) indicating that the cell volume fraction decreases across maturation time points in the chick tendon(McBride et al., 1985) and Kalson et al (2015) showed cell per unit volume reduction across late embryonic stages and up to 6 weeks post-natal in mouse tail tendon(Kalson et al., 2015). We verified that the number of cells per area in chick metatarsal flexor tendons decreased between E14 and E18 and E20 (Figure 2E), and furthermore that rigid paralysis led to an increase in the cell density at E20, to levels seen at E16 in normal development when immobilisation treatments began (Figure 4G). While only rigid paralysis affected cell density, it shows that it is responsive to the mechanical environment. Visualising cells within the developing tendon showed a number of striking changes in spatial organisation across developmental time. While cells were already aligned longitudinally at E14, the linear arrangements were narrow and close together with no clear distinction between the cytoplasm of linearly adjacent cells, apparently elongated and overlapping. The nuclei were extremely elongated with long axes running in the direction of the linear cellular arrangements. Over developmental time to E20, the nuclei became rounded, the linear columns further apart, with clear interfaces between the cytoplasm of linearly adjacent cells (Figure 2D and 3E). This indicates that cells are organised into discrete cuboidal, stacked arrangements by E20. This is reminiscent of observations from Serial Block-Face Scanning EM in mouse-tail tendons across a later time frame, up to 6 weeks post-natal(Kalson et al., 2015). Kalson et al (2015) proposed that the arrangement of cells stacked end to end is important for neighbouring cells to define sections of continuous vertical channels for assembly and growth of collagen fibrils; i.e. this arrangement is important for controlling collagen maturation(Kalson et al., 2015). We show that this is also the arrangement in chick tendons but is achieved at an earlier developmental time point. The increased distance between columns of cells up to E20 coincides with increased diameter of fibres(Kalson et al., 2015; Peterson et al., 2021) and is interpreted as creating space for more ECM deposition, and responsible for the reduction in cell volume to ECM ratio. Importantly, we found that both flaccid and rigid immobilisation caused a retardation in this progression toward the more mature arrangement, with rigid having a stronger effect than flaccid (Figure 4I). Therefore, one of the impacts of the dynamic mechanical environment generated by movement is the correct spatial organisation of tenocytes, appropriate for the maturation of the ECM.

A central objective of this study was to investigate YAP signalling as a candidate biological mechanism that integrates mechanical signals during the critical period of tendon maturation. It is a prime candidate mechanotransduction pathway to investigate due to is proven role in numerous biological contexts(Deng et al., 2016; Dupont et al., 2011; Shea et al., 2020; Wada et al., 2011), with emerging evidence of a potential role in tenogenesis in vitro(Li et al., 2021; Lu et al., 2023; Tao et al., 2021; Tomás et al., 2019; Xu et al., 2022). Here we showed the localisation of YAP and pYAP in the tendon across stages and the expression of YAP target genes revealed significant increases across the period of maturation, indicating that the pathway is active and modulated (Figure 5). Following rigid immobilisation, there was an increase in the levels of all three target genes assayed (*Ctgf, Cyr61 and Ankrd1*) together with increases in tendon regulator genes *Tnmd* and *Mkx*, with no increase in collagen genes and a decrease in *Scx*. Flaccid paralysis only led to a significant increase in Ankrd1 expression indicating that the activity of the pathway was more highly impacted by rigid paralysis (Figure 5E). It is possible that immobilisation from an earlier developmental might lead to greater disturbance but this study was designed to focus on the biomechanical transition in tendon properties in later development. An investigation of nuclear versus cytoplasmic YAP localisation did not prove possible in embryonic tendon tissue, due a number of architectural features of the tissue making detection and specific definition of localisation difficult. These included the high density of cells in parallel alignment and the shape of the cells, especially in immobilised tissue (Figure 4E). This is the first study to investigate YAP signalling in the embryonic tendon, with movement manipulation. Our data suggests that YAP pathway output is tightly controlled during normal development and is impacted by mechanical cues, most strongly in the constant force environment of rigid paralysis. It is somewhat surprising that there is an increase in target gene expression across development and yet paralysis leads to a further increase in expression levels rather than a decrease. This may be due to the constant static stimulation of the rigid state. The increase in Ankrd 1 expression in flaccid paralysis may be due to different regulator inputs for this particular gene.

A surprising finding in this study is the change in shape of tenocyte nuclei as development proceeds from E14 to E20, going from extremely elongated to more round (Figure 2F). It has been extensively reported that nuclei in adult tendons are elongated(Freedman et al., 2022), however tendons with more elongated nuclei are seen at 8 and 19 months compared to 1 month old rat achilles tendons(Freedman et al., 2022). Our finding is in contrast to that of the embryonic chick jaw-closing tendon, TmAM, which shows nuclear elongation as development proceeds(Korntner et al., 2022). This difference could be due to the distinct anatomical location of these tendons, and perhaps an accelerated differentiation progression is required for jaw tendon function at birth for feeding. It is possible that nuclear shape is reflecting the mechanoregulator environment: the nucleus is the stiffest organelle and is known to respond to mechanical forces((reviewed in Dahl et al., 2008)). Signalling from the ECM through integrins and focal adhesion kinase can lead to alteration in the shape of the nucleus. Interestingly it has been shown that altering the shape of the nucleus by plating cells on a micropatterned substrate affected collagen 1 synthesis(Thomas et al., 2002). These observations will be further explored by comparing nuclear shape in developing mouse tendons.

This study has provided insight into the structural, cellular and molecular changes in embryonic chick tendons during late embryonic development and has expanded understanding of the impact of immobilisation on tendon maturation. We have shown for the first time that YAP signalling is altered following immobilisation across the important period of rapid maturation of tendon mechanical properties. Future work will focus on manipulation of the signalling pathway to further test its specific role in tendon maturation. We also describe several aspects of cell organisation and collagen deposition within the tendon that change across this period and are sensitive to movement. Collagen fibre alignment and the organisation of cells into end-to-end stacked columns, with increasing distance between adjacent columns over time, similar to cellular organisation observed at later stages in the mouse tendon and postulated to be important in fibril growth, were also shown to be altered following immobilisation, with differential effects of rigid and flaccid paralysis. We conclude that tendon maturation requires force exerted by associated muscles, with specific aspects requiring a highly controlled level of dynamic stimulation. This study takes a developmental approach to understanding how tendons are constructed and will form the important basis for future work to engineer improved tensile load-bearing tissues. It illuminates the mechanistic basis of mechanoregulation during development as an important step in knowing what molecular cues and signals to provide during tissue construction *in vitro*.

## 6 Author Contributions

RR performed experiments, conceptualised experiments, analysed data, drafted the manuscript.

ETB performed experiments, analysed data, drafted the manuscript.

PM performed experiments, conceptualised experiments, analysed data, drafted the manuscript.

LS generated preliminary data, contributed to discussion of the findings.

GH generated data

ND contributed to discussion of the findings.

NB contributed to discussion of the findings.

HMC contributed to discussion of the findings.

SS, conceptualised experiments, contributed to discussion of the findings, drafted the manuscript.

## Acknowledgments

The authors are grateful to Natalie Jablonski, Luiza Holzl and Tom Murphy for their assistance with this research. Thanks to Simon Carroll and Tugdual Haffner in the Trinity Centre for Bioengineering for guidance on PLM and to Marcin Michal Baran for advice on the Li-Core Odyssey system. The research was funded by the Science Foundation Ireland US-Ireland R& D Partnership programme (Ref20.US/3718) to PM

