## supplementary figure for "Embryo movement is required for limb tendon maturation"

### *Supplementary Material*

#### Supplementary Figures

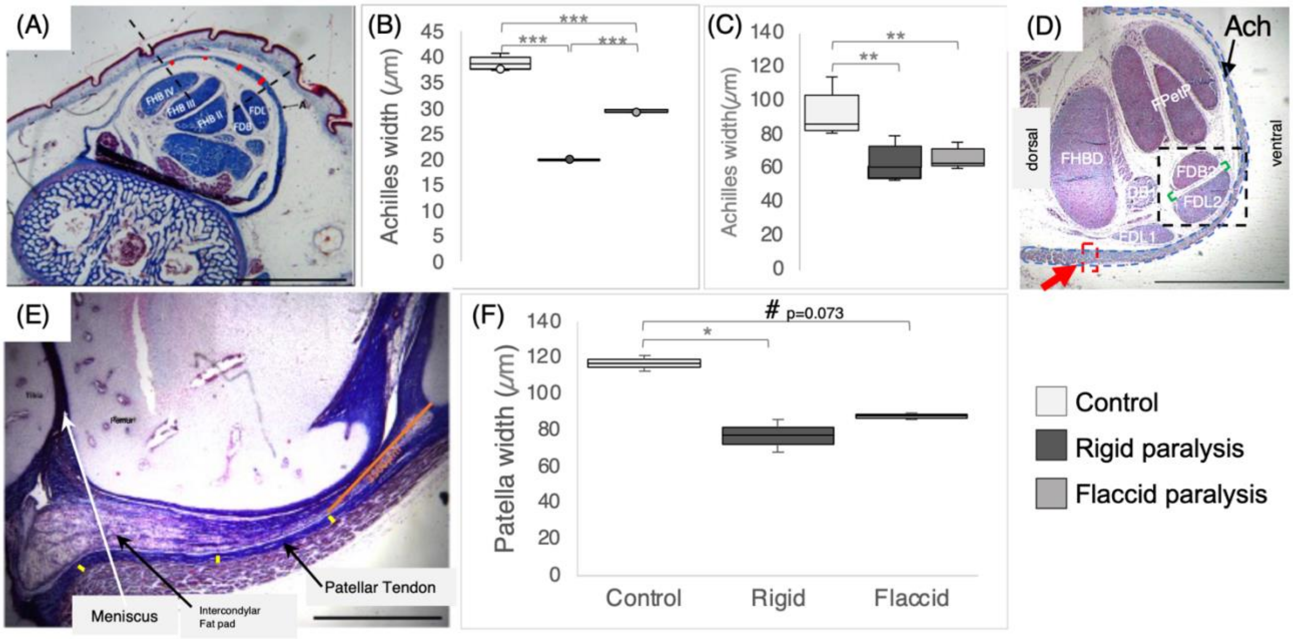


**Supplementary figure 1:** To confirm the finding of reduced Achilles tendon area under rigid and flaccid immobilisation (Figure 2), width measurements across consistent positions of the tendon were also taken.

<u>(A-D) The effect of immobilisation on the width of the Achilles tendon</u>. (A-B) Measurements averaged across 4 positions as indicated in A: Measurements were taken within the two dashed lines (placed consistently wrt flexor tendon morphology), at four equidistant locations indicated by red lines on the tendon (scale bar = 1000µm). (B) Box plot showing average measurements, control n=5, flaccid paralysis n=3, rigid paralysis n=3 (with 2-6 replicate measurements per specimen).

<u>(C-D) Width measurements taken at a consistent point in the Achilles tendon</u>. C, Box-plot of width of the Achilles tendon at the medial side of the Flexor Halluces Brevis Digits bundle, indicated by the red box and arrow in D; n=7 (control), n=6 (rigid), n=6 (flaccid). Abbreviations as in Figure 2. Taken together A-D show reduction in width across multiple positions within the Achilles tendon and confirm the reduction in size shown by area measurements.

<u>(E-F) The effect of rigid and flaccid paralysis on the width of the patellar tendon</u>. (E) the approach used to measure the width of the patellar tendon: Using midline longitudinal sections, landmarks were established to ensure 3 consistent measurements at different points along the ligament (shown by yellow lines); 1) 3500µm from the base of the patella (orange line); 2) in line with the knee joint; 3) halfway between 1 and 2. Scale bar 3000µm. F, The width of the patellar tendon is reduced with immobilisation. At least 7 measurements were taken for each specimen at E20; control n=2, flaccid paralysis n=2, rigid paralysis n=2.

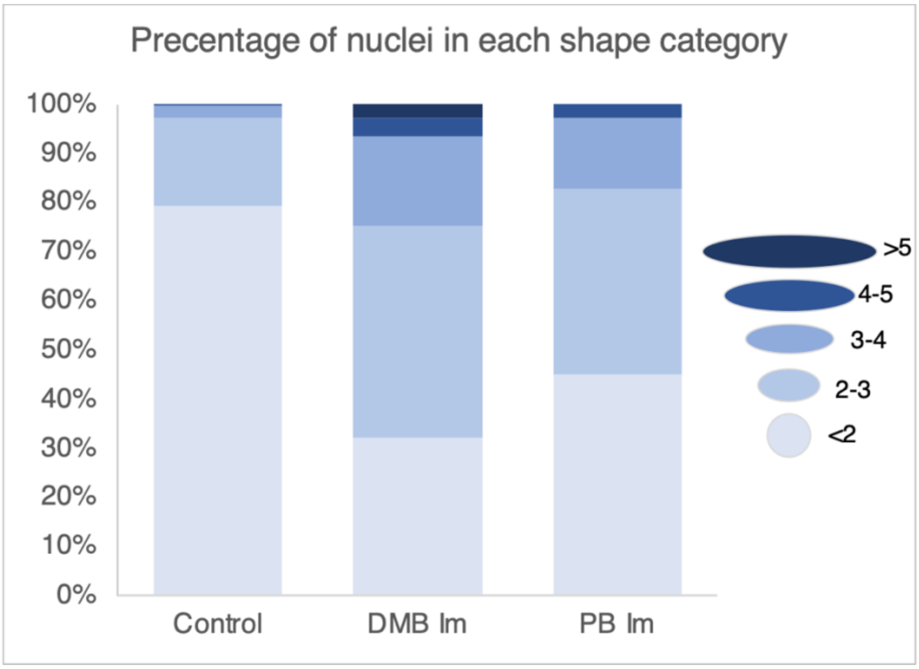


**Supplementary figure 2:** Nuclear shape distribution following paralysis. Histogram representing the proportion of nuclear aspect ratios following embryonic immobilisation, Rigid (DMB Im) and Flaccid (PB Im) compared to Control. Nuclear shapes are more elongated following immobilisation. Schematic of representative nuclei shapes in each category, with aspect ratio values indicated.

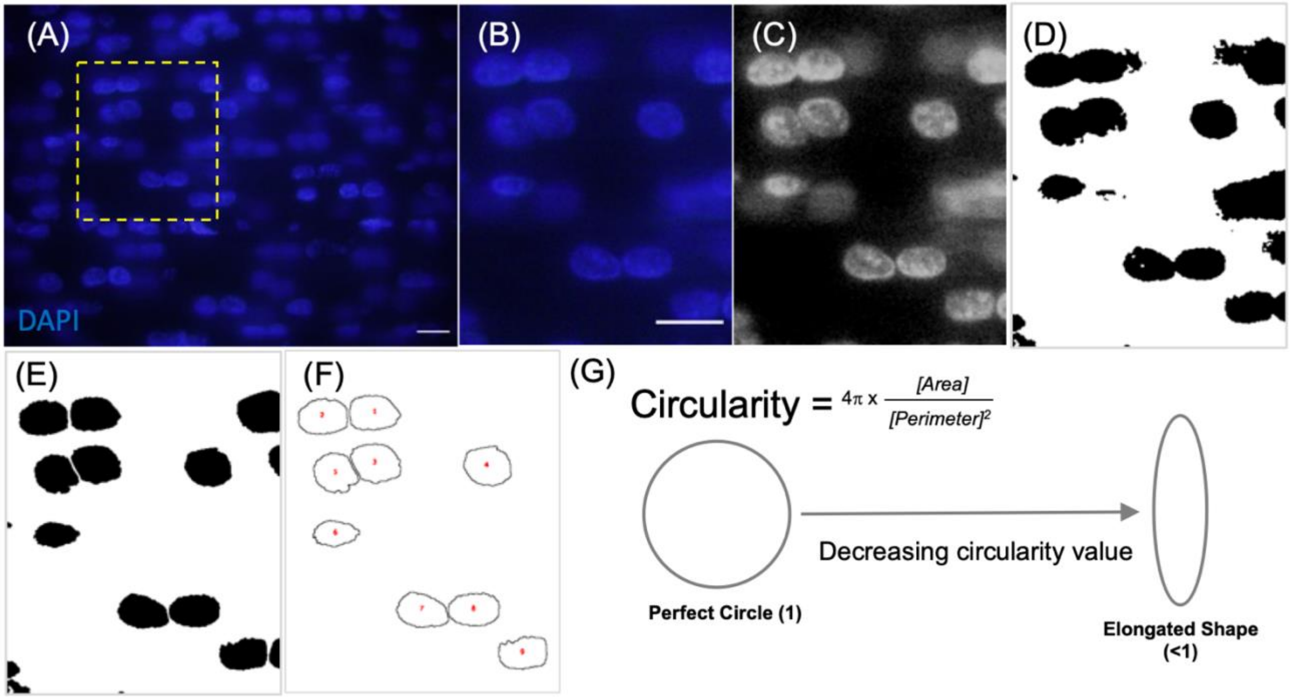

**Supplementary figure 3:** Methodological pipeline used to analyse the nuclear organisation pattern in embryonic tendons. (A) Original image of nuclei in the centre of longitudinal tendon section from tarsometatarsal region, (B) 4 x 5 $\mu$m rectangular cropped region of interest (ROI), (C) ROI split into component channels with the blue channel used for subsequent analysis. (D) conversion of blue channel into binarized black (nuclei) and white (space between nuclei) image, (E) noise was removed and any overlapping nuclei were separated, (F) encircling of each nuclei (minimum particle size of 0.05 $\mu m^2$) was performed with the analyze particles function. (G) formulae and illustration of how shape measurements were calculated for circularity, with values closer to 1 indicating a more circular shape. Scale bar; 1$\mu$m.
